# Modulation of the p75NTR during adolescent alcohol exposure prevents cholinergic neuronal atrophy and associated acetylcholine activity and behavioral dysfunction

**DOI:** 10.1101/2024.04.03.587970

**Authors:** Brian T. Kipp, M. Savage Lisa

## Abstract

Binge alcohol consumption during adolescence produces lasting deficits in learning and memory, while also increasing the susceptibility to substance use disorders. The adolescent intermittent ethanol (AIE) rodent model mimics human adolescent binge drinking and has identified the Nucleus Basalis Magnocellularis (NbM) as a key site of pathology. The NbM is a critical regulator of prefrontal cortical (PFC) cholinergic function and attention. The cholinergic phenotype is controlled pro/mature neurotrophin receptor activation. We sought to determine if p75NTR activity contributes to the loss of cholinergic phenotype in AIE by using a p75NTR modulator (LM11A-31) to inhibit prodegenerative signaling during ethanol exposure. Male and female rats underwent 5g/kg ethanol (AIE) or water (CON) exposure following 2-day-on 2-day-off cycles from PND 25-57. A subset of these groups also received a protective dose of LM11A-31 (50mg/kg) during adolescence. Rats were trained on a sustained attention task (SAT) while recording activity with a fluorescent acetylcholine indicator (AChGRAB 3.0). AIE produced learning deficits on the SAT, which were spared with LM11A-31. In addition, mPFC ACh activity was blunted by AIE, which LM11A-31 corrected. Investigation of NbM ChAT+ and TrkA+ neuronal expression found that AIE led to a reduction of ChAT+TrkA+ neurons, which again LM11A-31 protected. Taken together these findings demonstrate the p75NTR activity during AIE treatment is a key regulator of cholinergic degeneration.

## Introduction

Binge ethanol consumption during adolescence produces long lasting impairments in frontocortical-dependent cognition and executive function [1–3]. The observed executive dysfunction following adolescent binge drinking could be perpetuated by underlying deficits in attention. Of particular note, there has been limited investigations into the lasting effects that binge ethanol exposure during adolescence has on sustained attention performance. Attentional performance is critically modulated by cortically projecting forebrain cholinergic neurons [4, 5]. NbM cholinergic neurons from the basal forebrain send projections that terminate within the mPFC, orbitofrontal cortex (OFC), and anterior cingulate cortex. Reductions in this cholinergic population resulting from the neurotoxin saponin have been shown to coincide with impairments in sustained attention, which can be restored through administration of an M1 positive allosteric modulation [6]. Furthermore, optogenetic stimulation of cholinergic neurons within the NbM during cued trials of a SAT increased the number of correct responses, while stimulation during non-cued trials increased incorrect responses, and inhibition during cued trials increased the number of omissions [7]. Moreover, PFC increases in Acetylcholine (ACh) during a cued-appetitive response task coincide with behavioral indicators of cue detection that precede reward delivery [8]. These studies suggest that NbM cholinergic neurons drive the detection of environmental cues, and reductions in cortical ACh release contribute to impairments in attentional performance.

The cholinergic basal forebrain, in particular the NbM, actively participates in frontocortical-dependent tasks including in working memory, attentional processes, and response to reward and punishment [5, 7, 9–15]. In addition, recent work has identified an increase in NbM cholinergic activity during both decision making and reward approach in rats on radial arm maze task [16]. NbM cholinergic neurons respond transiently to both reward and punishment, and the magnitude of response appears to mirror the degree of uncertainty [12, 17]. Together, these suggests that basal forebrain cholinergic neurons participate in multiple aspects of cognition ranging from attention, learning, memory, as well as outcome valence. Thus, reductions in this neuronal population produce cognitive impairments across several cognitive domains. While we are beginning to understand the role the NbM plays in mediating behavior, it is uncertain the extent to which disease states, in particular developmental ethanol exposure, influences cholinergic signaling dynamics during complex cognitive tasks.

Adolescent ethanol exposure consistently leads to a persistent suppression of Choline Acetyltransferase (ChAT) expression within neurons [18–22]. Loss of the cholinergic phenotype in neurons from AIE leads to blunted ACh efflux within the frontal cortex, and cognitive impairments in both orbitofrontal cortical-dependent reversal learning, as well as medial prefrontal cortical-dependent cognitive flexibility [20, 23–25]. Although we have previously detected a significant reduction in PFC and OFC ACh efflux through in vivo microdialysis during a spontaneous alternation task, the temporal resolution of this method of detection is limited to the accrual of ACh release over several minutes and not appropriate for analysis of discrete ACh signaling dynamics to microbehaviors (choice decision, reward approach, cue detection, etc.). The recent development of an ACh sensor (GRABACh 3.0) is an ideal match for physiology assessments during cognitive testing as it permits the measurement of ACh activity on the order of seconds with in vivo fiber photometry [26–28].

Although we have consistently identified a loss of ChAT+ neurons in the basal forebrain following AIE, the upstream mechanisms that facilitate this degeneration remain elusive. A growing body of literature implicates the p75NTR, expressed by cholinergic neurons, in mediating disease states in response to proneurotrophin binding. In AD, loss of cholinergic neurons has been proposed to manifest following reductions in the ratio of Trk receptors to p75NTR [29, 30]. LM11A-31, a p75NTR inhibitor, that prevents proneurotrophin-p75NTR interaction by blocking the ligand binding site [31–35], is a treatment strategy that can determine if p75NTR drives the loss of cholinergic phenotype with AIE. Previous studies examining the effectiveness of LM11A-31 has been found to be effective in preventing/treating age-related cholinergic neurodegeneration [33, 36, 37]. AD models have demonstrated that modulating p75NTR is an effective means of preserving CBF, and LM11A-31 has not only prevented AD-associated CBF pathology, and has also been shown to reverse previous damage already produced by amyloid beta accumulation [32, 33, 35, 37]. Although there are limited examinations in the use of LM11A-31 during developmental timepoints, or with ethanol, this compound has been administered to Sprague Dawley rats at post-natal day 35, daily for two weeks without gross abnormalities [38], and has previously been used in an adult ethanol model to attenuate excessive alcohol consumption [31].

The goal of the proposed study is two-fold: First, is to investigate discrete AIE-associated deficits in cortical cholinergic activity during a sustained attention task. Secondly, we aim to elucidate whether the p75NTR mediates the loss of cholinergic phenotype- and more importantly, if modulating the p75NTR during developmental ethanol exposure conserves the cholinergic phenotype and protects cognitive performance in adulthood.

## Materials and Methods

### 2.1 Subjects

Eighty male and female Sprague Dawley rats that were bred from dams received from Envigo (Indiannapolis, IN) at Binghamton University were used generate 10 rats per sex per treatment condition. Rats were randomly assigned treatment conditions with no more than 1 subjects per sex from each litter. On PND21 male and female rats were weaned and double/tripled same-sex housed in a temperature and humidity-controlled vivarium under a 12-hour light-dark cycle (0700 – 1900) at Binghamton University. Rats were provided with ad libitum access to food chow (Purina Lab Diet 5012) and water and were provided with environmental enrichment in the form of nesting packets and wooden chew blocks.

### 2.2 Treatment

Rats were exposed to one of four treatment conditions (∼10 per sex/treatment/drug exposure). AIE-treated rats received intragastric gavages of 5g/kg 20% EtOH and control-treated rats (CON) received volume matched equivalents of water following a two-day-on two-day-off cycles from post-natal day 25-57 (See Figure 1 for a timeline of treatment and behavioral testing). The third treatment group, AIE-LM, received identical ethanol treatment as AIE, however, this group also received 50mg/kg of the p75NTR modulator LM11A-31 (MedChemExpress, Monmouth Junction, NJ, USA) through intragastric gavage 30 minutes before, and 8 hours following each water or EtOH gavage. The last treatment condition CON-LM underwent the same treatment schedule as CON treated animals; however, this group also received 50mg/kg LM11A-31 30 minutes before and 8 hours following each gavage. In addition to water and ethanol administered, rats that did not receive LM11A-31 received an additional intubation of water (vehicle; V) to match volume of LM11A-31 administered. One hour after the 8^th^ gavage, BECs were collected from tail veins of all treatment conditions and were measured (GM7 Analyzer, Analox, London, UK).

**Figure 1:**
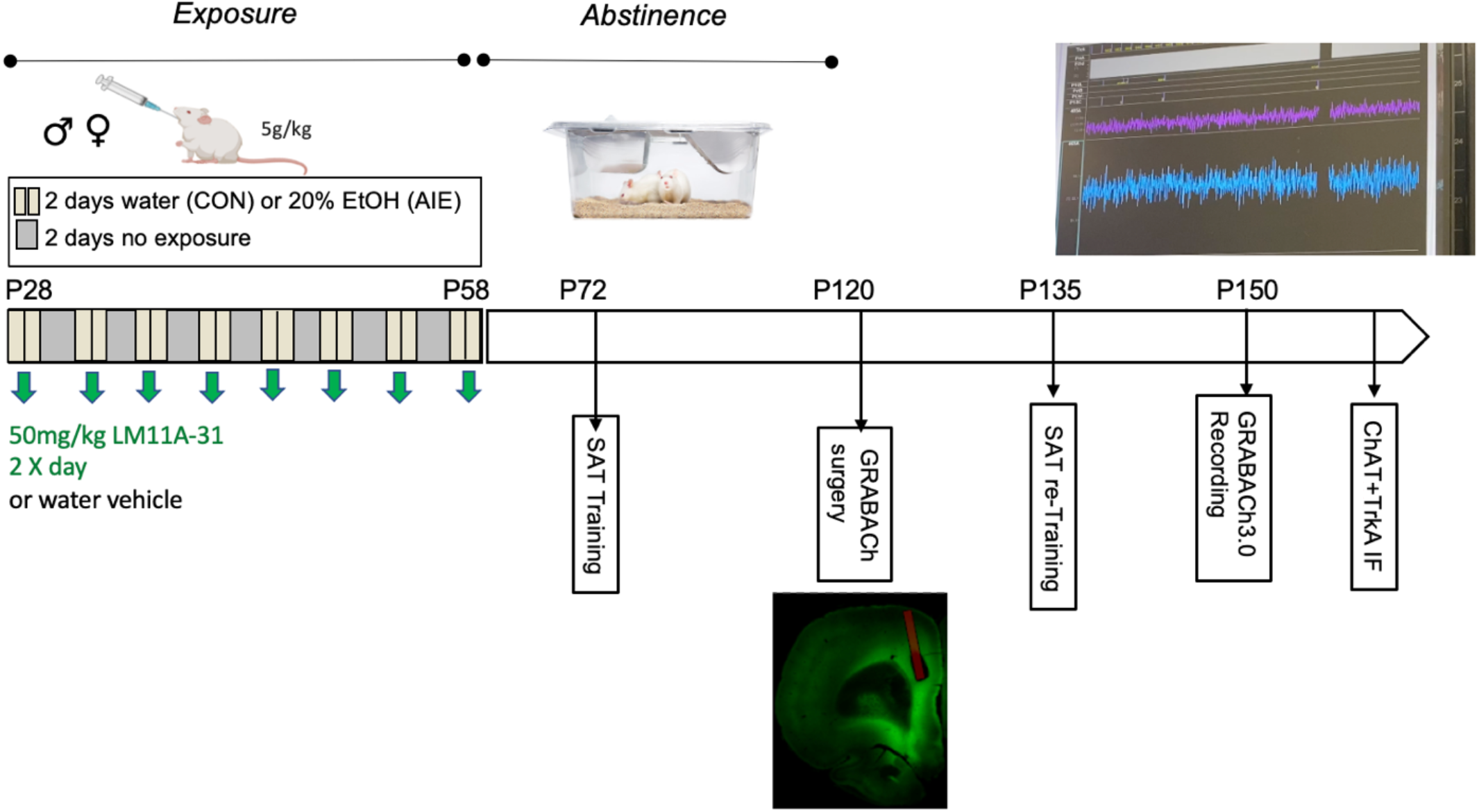
Experiment 2 Treatment Timeline. Groups underwent the previously described AIE or CON treatments. A Separate AIE+LM11A-31 and CON+LM11A-31 treatment groups received IG gavage of 50mg/kg LM11A-31, 30 minutes before and 8 hours following each gavage. Animals were trained on SAT 2 weeks following the end of AIE treatment. Following mastery of SAT, subjects underwent ACh GRAB 3.0 viral infusion into the mPFC with fiber optic cannula 2 weeks after the end of AIE. 3 weeks following viral infusion, and SAT re-training, groups underwent operant pretraining and behavioral testing on the SAT where mPFC ACh activity will be recorded through in vivo fiber photometry.

### 2.3 Operant Sustained Attention task

Operant chambers (30cm x 33 cm x 23 cm; Med Associates Inc., St. Albans, VT, USA) located within a sound attenuating shells (59 cm x 55 cm x 36 cm), interfaced with MED-PC IV (Med Associates Inc.) and Synapse recording software (Tucker Davis Technology, Alachua, FL, USA) were used for operant training and fiber photometry recording. Each operant chamber contained two retractable levers situated on the left and right of a food trough through which a single food pellet (Rodent Purified Dustless Precision Pellet; Bio-Serve, Flemington, NJ) was dispensed upon correct responses. The operant chambers also contained a single cue light located directly above the food trough and a house light positioned on the opposite wall of the chamber. Behavioral inputs from the operant chamber were timestamped in Synapse via TTL pulses using the digital input and output ports from ICON4 system (Tucker Davis Technologies) that interfaced directly with the fiber photometry recording system (RZ10x Processor; Tucker Davis Technology). All rats were food restricted to 85% of their baseline free-feed weight over the course of one week and were maintained at this restriction, determined by a growth curve, throughout behavioral testing. In addition, rats were handled on the week leading up to operant training.

Two weeks after the end of AIE treatment rats underwent pretraining and an operant SAT task following a Fixed Ratio – 1 (FR-1) schedule. During the pretraining phase, rats were conditioned to lever press the left and right lever for 30 minutes across two consecutive days. On the third day of pretraining rats began retractable lever training. Retractable lever training is comprised of 90 trials with the goal of training rats to respond within 10 seconds of lever presentation. This training also occurred on a FR-1 schedule. Rats were required to make fewer than 5 trial omissions within 90 trials for two consecutive days.

Following completion of pretraining rats were trained on the SAT pretraining (pSAT). pSAT is a simple discrimination task composed of 81 cued and 81 non-cued trials (Figure 2). Rats are trained to detect the illumination of a central cue light of varying duration (500ms, 50ms, and 25ms) that occur 2 seconds prior to lever presentation. On cued trials, rats were rewarded for pressing the left lever (Hit, h) and a right lever press was not rewarded (Miss).

**Figure 2:**
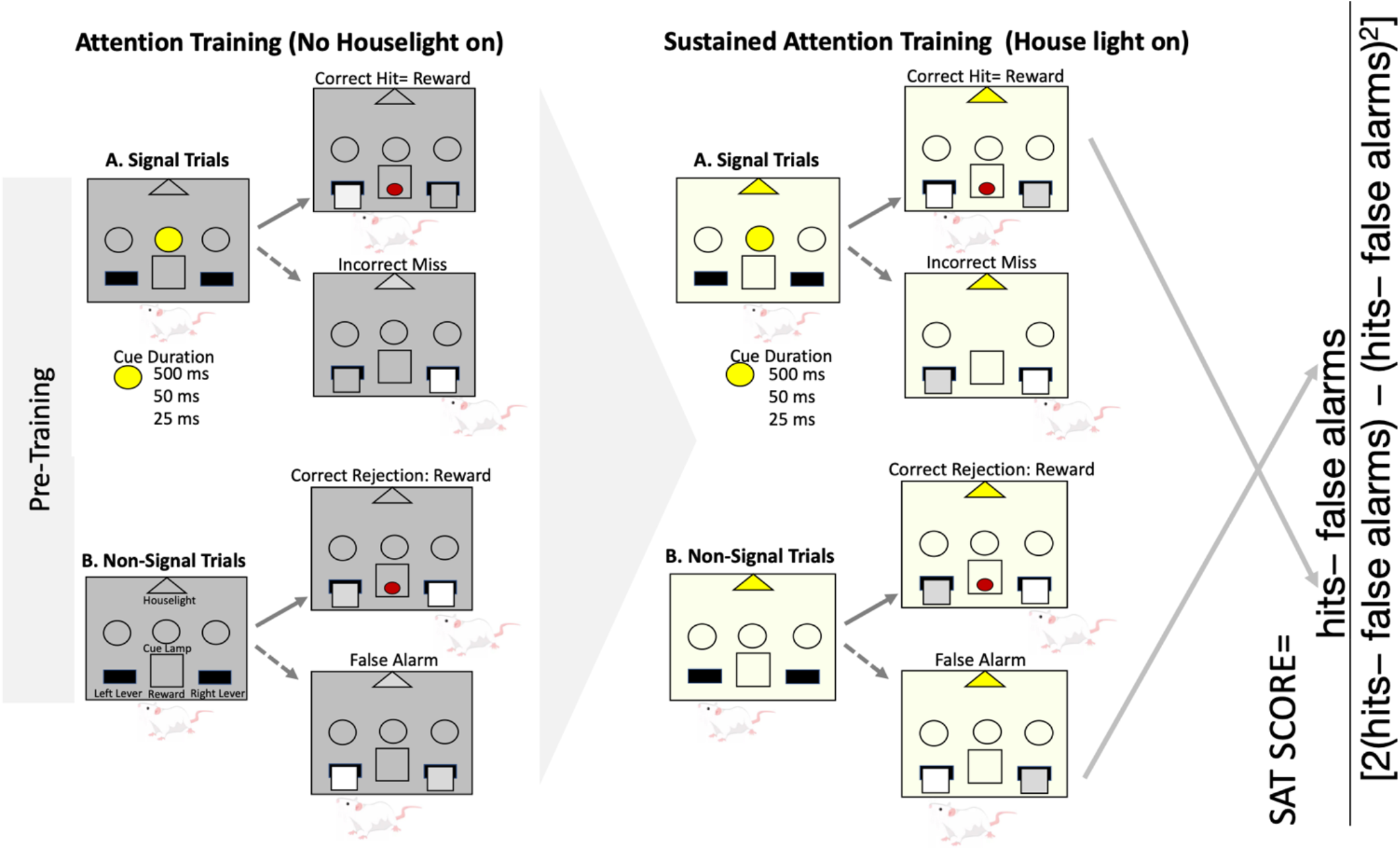
Sustained Attention Task and SAT Score. Depiction of the SAT pretraining, SAT Tasks, and formula for the development of the SAT score. Animals underwent 162 trials per session, 81 of which were non-cue trials while 27 cue trials were utilized with 500ms, 50ms, and 25ms cue duration (total of 81 cue trials). Attention training (pSAT) occurred in the absence of the house light, but after mastery of the pSAT condition, the house light was introduced. The SAT score was calculated using the above formula consisting of hits at each cue duration and total false alarms.

Conversely, on non-cued trials, a left lever press was not rewarded (False Alarm, f), whereas right lever presses were rewarded (Correct Rejection). pSAT took place without general illumination provided by the house light. Mastery of the task was assumed when rats had with 3 consecutive sessions with > 70% Hits at the 500ms cue duration and > 70% Correct Rejections. Following this criterion, rats were moved onto the SAT task that had under general illumination of the house light. The inclusion of the house light requires the rats to be vigilant to the cue [39]. The same task contingencies that occurred under pSAT applied to the SAT. Four female rats were unable to meet the passing criteria of the SAT task and were excluded from all measures (two females from the CON group, one female AIE, and one female AIE-LM).

Since this task utilizes two different types of stimuli to be discriminated, signal detection theory measures were calculated to thoroughly evaluate performance on this task [6, 40–43]. First, the SAT score was calculated using the following formula (h-f)/(2(h+f)-(h+f)^2^ ), generating a value ranging from 1.0 (all responses on cued trials were hits, and all responses on non-cued trials were correct rejections) to -1.0 (all responses on cued trials were misses, and all responses on non-cued trials were false alarms). It was determined that chance performance on this measure is 0 ± 0.17 [6]. Two additional measures derived from signal detection theory were utilized to determine both perceptual sensitivity (A’), as well as response bias (B”) [41, 43, 44] A’ is a subject specific measure of strength of signal relative to noise and is unaffected by response bias. A’ was calculated as [0.5+((h-f) (1+h-f)/4h(1-f))], producing a value ranging from 0.5 (subject cannot distinguish signal from background) to 1 (subject always registered a hit and never registered a false alarm). B”, on the other hand, is a measure of response bias calculated as ((h(1-h)-f(1-f)/(h(1-h) +f(1-f)), generating values ranging from 1.0 (respond that the cue was never present) and -1.0 (respond that the cue was always present).

### 2.4 ACh Grab 3.0 Viral infusion Surgery and ACh 3.0 Recording

One week following mastery of the SAT task, subjects underwent stereotaxic surgery in preparation for in vivo fiber photometry recording. Rats were administered a mixture of Dexmedetomidine (Dexdomitor, Zoetis, Kalamazoo, MI; 0.04mg/kg) and Ketamine (Ketaset, Zoetis, Kalamazoo, MI; 85mg/kg) intraperitoneally, for anesthesia. A genetically encoded calcium indicator for the detection of ACh activity (1 μL, AAV9-hSyn-ACh3.0; AddGene; Watertown, MA) was infused into the medial prefrontal cortex (AP(2.7mm); ML (+/-0.7mm); DV(-3.0mm) [45]), prior to fiber optic cannula placement (MFC_400/430-0.48_5mm_ MF1.25_ FLT, Doric Lenses, Quebec, Canada), at the same AP and ML coordinates, but 0.1mm dorsal to the site of viral infusion. At the end of surgery, each subject received an intraperitoneal injection of Antisedan (Atipamezole HCl, 5g/kg) to reverse the sedation and were returned to home cages to recovery. Subcutaneous injections of carprofen (5g/kg, Zoetis, Kalamazoo, MI) were used as an analgesic prior to surgery, 24 hours post-surgery, and 48 hours post-surgery. Rats were then allowed to recover for 3 weeks prior to behavioral testing and fiber photometry.

Following viral infusion and cannulation, animals were retrained to the pSAT and SAT tasks with the stated criteria. On the session following the third day of passing the SAT, a one-meter-long fiber optic patch cord (MFP_400/430/1100-0.57_1m_FC-MF1.25_LAF, Doric Lenses) was attached to the fiber optic cannula, and rats were left to habituate in the operant chamber for 30 minutes before collecting a baseline signal of AChGRAB 3.0. Following baseline collection, the SAT task was started with recording of AChGRAB 3.0. The ACh indicator was excited with a ∼490nm signal and a second ∼405nm signal was used as a control. While the signal collected from the ∼490nm channel reflects the ACh binding to a GFP conjugated nonfunctional M3 ACh receptor, the ∼405 isosbestic control channel is used to correct for motion artifact and background fluorescence. Signals emitted from AChGRAB 3.0 were collected with the signal processor (RZ10x Processor) running Synapse software (Tucker Davis Technology) through the same fiber optic patch cord. Behavioral inputs were timestamped onto the collected 490nm and 405nm channels via TTL pulses that marked cue light presentation, lever extension, and lever pressing, and reward delivery. We were unable to collect fiber photometry data from 6 subjects due to loss of head cap prior to meeting passing criteria (one male CON, one female CON, two female AIE, two female CON-LM).

### 2.5 ACh 3.0 Data Analysis

All analyses for AChGRAB 3.0 activity were conducted in Spyder 5.0.3 running a kernel of Python 3.4. Signal collected from the 490nm channel was transformed to ΔF/F by subtracting fluorescence detected in the 490nm channel from the 405nm channel before dividing the result by the signal in the 405nm channel. Peri event histograms were produced by measuring the ΔF/F 2 seconds before and after each time-stamped behavioral event. Peri-event histograms are then plotted as a normalized z-score of the entire recording session. The Z-score of rats belonging to the same treatment group were then averaged at each behavioral event. To account for photobleaching over the course of the SAT task, only the first 20 minutes of AChGRAB 3.0 activity was used for analyses. From the generated Z score, peak Z-scores pertaining to behavioral events in cued trials (Cue presentation preceding Hits, Cue presentation preceding Misses, Hits and Misses) at each time interval were measured alongside the peak Z-score during Correct Rejections and False alarms of non-cued trials. Area under the curve was also calculated using sklearn.metrics.auc to measure the total fluorescent output recorded during each task epoch relative to the within trial baseline period.

### 2.6 Tissue Collection

One week after the end of operant training, rats were sacrificed (Fatal-Plus, Vortech Pharmaceuticals, Dearborn, MI), and transcardially perfused (Master Easy-Load Console Drive, 7518-00; Cole-Palmer Instruments) with cold phosphate buffer saline and cold 4% paraformaldehyde (PFA; Electron Microscopy Services, Hatfield, PA). Tissue was then post fixed for 24 hours in 4% PFA before transferring to a 30% sucrose solution in 0.1M PBS and stored at 4°C. Brains were sliced coronally on a freezing sliding microtome (Sm2000r; Lecia Biosystems, Wetzler, Germany) at 40μm and then stored at -20°C in cryoprotectant (62.8mg NaH_2_HPO_4_, 160mL dH_2_O, 120mL ethylene glycol, and 120mL glycerol). In order to confirm AChGRAB 3.0 viral and fiber optic cannula placement, mPFC tissue sections were immediately mounted onto gelatinized slides and cover slipped with Prolong Glass Antifade (Thermo Scientific, Waltham, MA) and imaged at 10x using an Olympus VS200 Slide Scanner.

### 2.7 ChAT and TrkA Immunofluorescence

Four sections per subject (approximately -0.72, -1.08, -1.44, and -1.8AP) were used to investigate treatment effects of AIE and LM11A-31 on the expression of ChAT+/TrkA-, and ChAT+/TrkA+ cell types within the NbM. Free floating sections were washed in 0.1M Tris Buffered Saline (TBS), followed by a 30-minute quench process in 0.3% H_2_O_2_ in 0.1M TBS. NbM sections were again washed in 0.1M TBS before undergoing an antigen retrieval step where tissues were incubated in a sodium citrate buffer at 80°C for 30 minutes. Sections were again washed in 0.1M TBS prior to being placed in a blocking solution (4% normal donkey serum, 0.1% Triton X-100) for one hour. Tissues were then transferred to a ChAT primary and TrkA primary antibody solution with blocking buffer (1:200 dilution goat anti-ChAT polyclonal AB143, Millipore EMD, Billerica, MA; 1:200 rabbit anti-TrkA polyclonal 06-574, Millipore EMD, Billerica, MA) and incubated overnight at 4°C. Following 3x 5-minute washes in 0.1M TBS, NBM sections were then incubated in blocking buffer with secondary antibody (1:300 dilution Alexa fluor 488 Donkey anti Rabbit, Thermoscientific, Waltham, MA; 1:400 dilution Alexa Fluor 680 Donkey anti Goat, Jackson Immuno Research, West Grove, PA) for 2 hours. Afterwards, sections were then washed in 0.1M TBS before cover slipped with Prolong Glass Antifade (Thermoscientific, Waltham, MA)

### 2.8 Microscopy and ChAT/TrkA Cell Counting

Slides were randomly coded and imaged at 20x using an Olympus VS200 Slide Scanner. Images were exported to FIJI through the Olympus plugin. NIH Image J cell counting tool was used to quantify the total number of ChAT+/TrkA-, and ChAT+/TrkA+ cholinergic neuron phenotypes in one hemisphere across all NbM sections. Since only one hemisphere was counted, the total number of ChAT+TrkA+ and ChAT+TrkA-cells were doubled.

### 2.9 Statistical Analyses

For the measurement of change in body weight over the course of treatment, a repeated measures ANOVA was used with between-subjects factors of treatment condition (AIE and CON) and drug (LM11A-31 and Vehicle), the within subject factor was day of gavage. Male and female analyses for change in body weight over the course of treatment were run separately. BECs were measured using a 2-way ANOVA (Drug, Sex) and AIE and CON were run separately, such that the analysis reflects AIE comparison to AIE-LM and CON comparison to CONLM. Operant data for the pSAT and SAT tasks prior to, and post-surgery were run using a 3-way ANOVA (Treatment, Drug, and Sex) with Fishers LSD post hoc analyses used to examine group differences when the omnibus F was significant. Since analysis of the fiber photometry data revealed no significant main effects of Sex in the 3-way ANOVA (all p’s. > 0.05), sex as factor was collapsed, and data were analyzed as a 2-way ANOVA with Fishers LSD post hoc analyses. Lastly, NbM ChAT+TrkA+, and ChAT+TrkA-data were analyzed using a 3-way ANOVA (Sex, Treatment, Drug) with Fishers LSD post hoc analysis for group differences. All statistical analyses were conducted in Prism 9.

## Results

### 2.10 Treatment Growth Curves and BECs

#### Male Growth Curve

When analyzing change in body weight of males over the course of AIE treatment, it was found that Mauchly’s test of sphericity was violated (p < 0.001), therefore a Greenhouse-Geisser correction was applied for within-subject analyses. Over the course of treatment, all males significantly increased in body weight from PND 25-57 (F(1.85,64.74) = 4453.924, p < 0.001; Fig 3A). However, CON, water treated, rats had a greater increase in body weight during treatment than AIE males (F(1.85, 64.74) = 11.724, p < 0.001; Fig 3A). Treatment with LM11A-31 did not influence change in body weight during development (p > 0.05).

**Figure 3:**
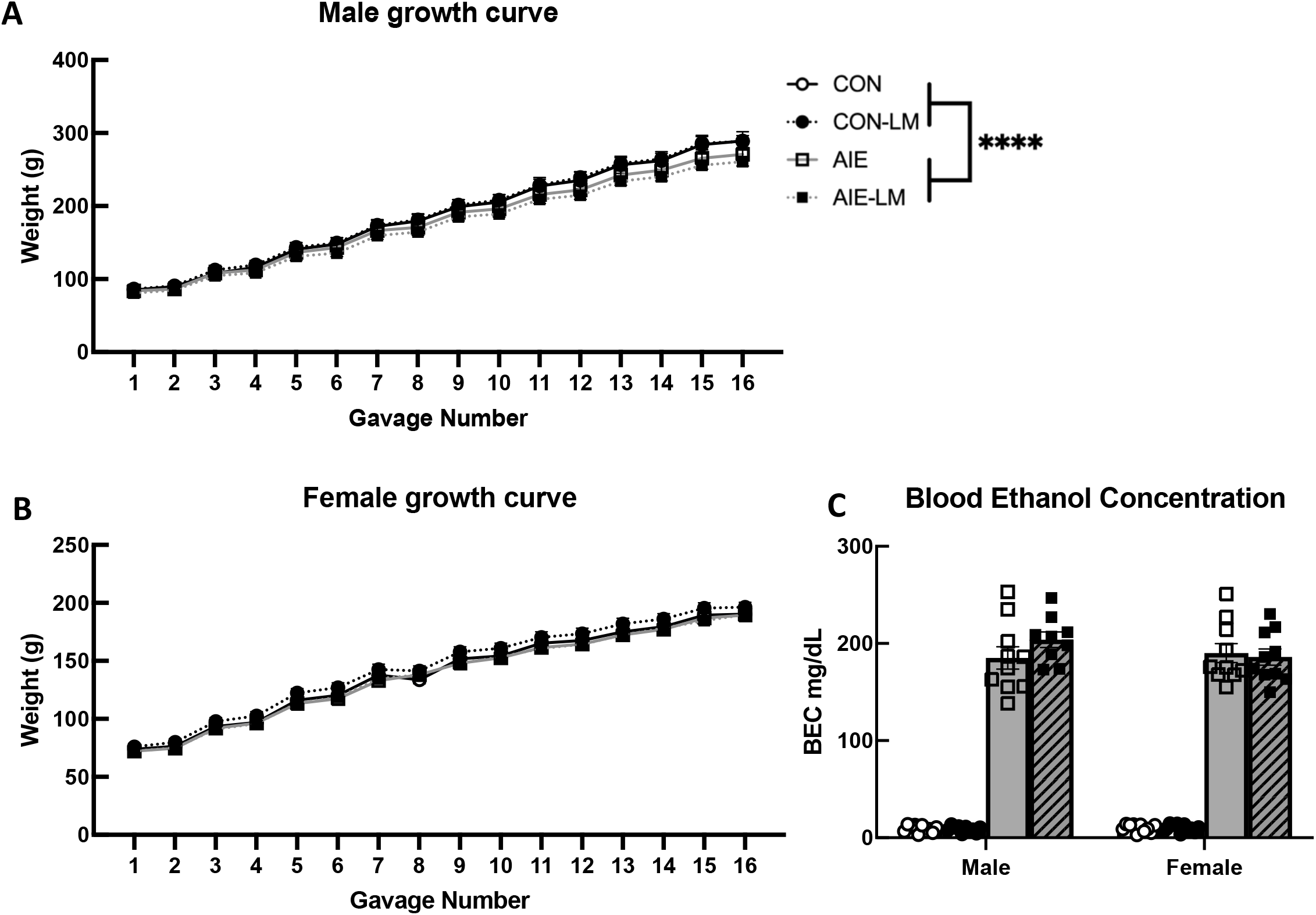
Growth Curves. Graphs depicting the change in body weight of male and female Sprague Dawley rats over the course of AIE or CON treatment with or without LM11A-31 administration. All subjects significantly increased weight gain over the course of treatment. In males, CON-V and CON-LM males gained more weight than AIE-V and AIE-LM males (**A**). Similarly, female weight gain over the course of treatment did not significantly differ across CON and AIE treatment conditions, or with the administration of LM11A-31. (**B**). Blood ethanol concentration measured from tail bloods collected 1 hour following the 8^th^ gavage. LM11A-31 treatment did not significantly alter BECs in CON or AIE treated animals during treatment in adolescence (**C**) Data represent group average ±SEM. **** Indicates p < 0.0001.

#### Female Growth Curve

A Greenhouse-Geisser correction was also applied to within subjects analyses for female growth curve data as Mauchly’s test of sphericity was violated (p < 0.001). Similar to males, female weight also significantly increased during treatment from PND 25-57 (F(2.74, 98793) = 2218.26, p < 0.0001) and LM11A-31 did not alter growth trajectory in adolescence (p > 0.05). Unlike the males, females weight change was not affected by AIE (p > 0.05; Figure 3B).

#### Blood Ethanol Concentrations

To determine if LM11A-31 affected blood ethanol concentrations (BEC) during AIE treatment, tail bloods were collected following the 8^th^ gavage revealing that CON-LM did not differ from CON-V treated rats (p = 0.97), nor were sex differences present (p = 0.38; Figure 3C). The comparison of AIE-LM and AIE animals revealed that LM11A-31 did not affect BEC during AIE treatment (p = 0.44), and no differences were observed between males and females (p = 0.50; Figure 4). In both cases, no significant Sex or Sex X Drug interactions were detected (p’s > 0.05).

**Figure 4:**
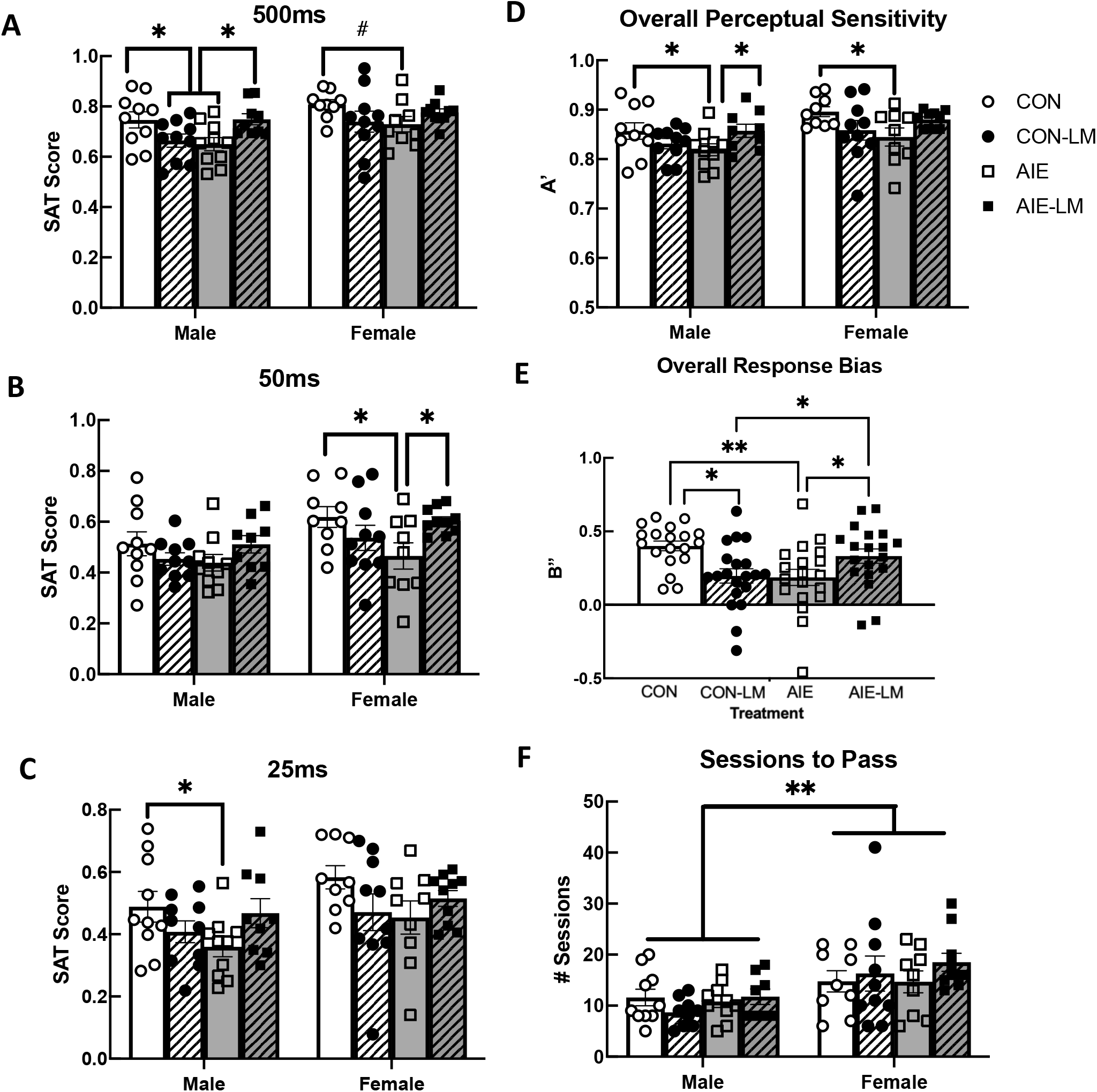
pSAT performance. SAT score and Signal Detection Theory measures on the SAT task in the absence of the house light (pSAT). Data represent group mean ±SEM. AIE-V had lower SAT scores, which was recovered in AIE-LM males. However, LM11-31A treatment in CON male rats had the opposite effect, it impaired performane. AIE-V females performed worse than CON-V females on 500ms trials, an effect not seen in rats that received LM11-31A (**A**). Under the 50ms cue duration, AIE-V females performed significantly worse than CON-V and AIE-LM females; however, no group differences were found in male rats (**B**). At the 25ms cue duration, only male AIE-V rats performed worse than their CON-V counter parts, an effect not seen in LM11-31A treated rats (**C**). Measures of perceptual sensitivity in AIE-V treated male and female rats were lower than male and female CON-V rats, which was not seen when LM11-31A was given (**D**). Regardless of sex, AIE-V and CON-LM treated rats had a more liberal response bias compared to CON-V and AIE-LM rats (**E**). Lastly, females overall, took significantly more sessions to complete pSAT than males, regardless of treatment (**F**). * Indicates p < 0.05; ** indicates p < 0.01

### 2.11 SAT pretraining (pSAT)

The number of sessions required to pass pSAT task was investigated for group differences, revealing a significant main effect of Sex (F(1,36) = 12.4, p = 0.0012), where on average, females (16.13, SEM = 1.21) took significantly more sessions to reach criterion than males (11.64, SEM = 0.94). Since males and females differed significantly in the number of sessions to pass pSAT, separate analyses were conducted to determine if Treatment and/or Drug effects were present in males and females. In both males and females, the number of sessions required to pass pSAT was unaffected by Treatment, Drug, or the interaction of the two factors (all p’s>0.15).

To determine if cue duration significantly impacted performance on the pSAT task, a within subjects ANOVA was conducted on SAT scores across the 500ms, 50ms, and 25ms cue durations (see Figure 4ABC). Sphericity was found to have been violated (p<0.001), therefore a Greenhouse Geisser correction was applied. As the cue duration shortened, SAT scores were reduced regardless of treatment (F(1.56, 98.43) = 374.97, p < 0.0001). Group differences were also investigated during the pSAT task at each cue duration to identify underlying group differences in attentional performance. When examining the 500ms cue duration, a significant main effect of Sex was observed (F(1,36) = 8.47, p = 0.006), where females performed significantly better than males (0.70, SEM = 0.01). Therefore, males and females were run in separate analysis. In males, a significant Treatment X Drug interaction was detected (F(1,35) = 11.1, p = 0.002). Post hoc analyses revealed that male CON-V rats had higher SAT scores compared to AIE-V male rats (p = 0.015) and CON-LM males (p = 0.03). Additionally, male AIE-LM treated animals performed better at the 500ms cue duration that AIE-V treated animals (p = 0.015). When examining pSAT performance in females a trending Treatment X Drug interaction was observed (F(1,34) = 3.99, p = 0.053). Post hoc test revealed a trending difference where female AIE rats had lower SAT scores than CON females.

Similar to the 500ms cue duration, females had significantly higher SAT scores at the 50ms cue duration than males (F(1,36) = 7.484, p = 0.0096), therefore males and females were run in separate analyses. At the 50ms cue duration in males, no Treatment (p = 0.86), Drug (p = 0.81), or Treatment X Drug interactions were observed (p = 0.07). However in females, a significant Treatment X Drug interaction was detected (F(1,33) = 11.03; p = 0.012). Post hoc tests revealed that AIE-V treated females had lower SAT scores compared to CON females and AIE-LM treated females (p = 0.025).

Under the 25ms cue duration females outperformed males on the pSAT task (F(1,36) = 4.88; p = 0.033). Since males and females differed significantly at the 25ms cue duration, males and females were run separately. In males, at the 25ms cue duration, a significant Treatment X Drug interaction was detected (F(1,35) = 5.22; p = 0.028). Post hoc tests determined that male CON rats had higher SAT scores compared to male AIE rats (p = 0.031). In females, no significant differences were observed across treatment conditions (p = 0.36) or Drug (p = 0.58), and no Treatment X Drug interaction was found (F(1,34) = 3.57, p = 0.067).

Measures of perceptual sensitivity and response bias allow for examination of attentional performance without influence of the behavioral strategy animals used to pass the task (see Figure 4CDE). It was discovered that males and females significantly differed in perceptual sensitivity (F(1,36) = 8.357, p = 0.0095), where females had higher perceptual sensitivity scores compared to males. Since males and females differed significantly in A’, males and females were separated prior to subsequent analyses of treatment of drug effects. In males, a significant Treatment X Drug effect was observed (F(1,35) = 7.73, p = 0.0087). Post hoc analyses revealed that male CON and AIE-LM (p = 0.048) treated rats had significantly higher A’ scores compared to AIE-V treated males. In females a significant Treatment X Drug interaction was detected (F(1,34) = 5.17; p = 0.029). Further post hoc tests determined that female CON rats scored higher on perceptual sensitivity than AIE treated females (p = 0.021).

Regarding response bias, a main effect of Sex was not detected (p = 0.065), therefore, sex was collapsed across groups for further analyses. A significant Treatment X Drug interaction was detected (F(1,73) = 13.59, p = 0.0004). Post hoc analyses revealed that CON-V treated animals utilized a more conservative response strategy compared to CON-LM treated animals (p = 0.03), and AIE-V treated rats (p = 0.002). In addition, AIE-LM rats demonstrated a more conservative response strategy than AIE-V treated rats (p = 0.03) and CON-LM treated rats (p = 0.045).

Performance on pSAT was examined for group differences in the latency to lever press and latency to collect reward. Neither Treatment, Drug, nor their interaction, were found to have affected latency to lever press on pSAT. Males and females did not differ significantly in the latency to lever press on 500ms trials (p = 0.10). For the 50ms cue duration, a significant main effect of Sex was detected (F(1,68) = 6.54, p = 0.012), where females (370ms, SEM = 26.2) were significantly slower to lever press compared to males (310ms, SEM = 10.1). Similarly, a significant main effect of Sex was also detected at the 25ms cue duration (F(1,68) = 6.56, p = 0.012) where females (370ms, SEM = 27.8) were significantly slower to respond to lever presentation compared to males (310ms, SEM = 11.0). During non-cue trials, latency to lever press was unaffected by any variable (all p’s>0.22).

Latency to collect reward was also measured following correct response to trials. On 500ms trials, a significant Treatment X Sex interaction was present (F(1,30) = 5.01; p = 0.03), female CON-V and CON-LM females (390ms, SEM = 36.2) collected the reward more quickly than CON-V and CON-LM males (580ms, SEM = 129.9). In males, latency to collect reward was unaffected by AIE, LM11A-31, or the interaction (all p’s >0.15). Similarly in females, neither Treatment, Drug or the Treatment X Drug interaction (all p’s > 0.50) affected latency to collect reward. At the 50ms cue duration, a Sex X Treatment interaction was found (F(1,31) = 7.54; p = 0.01). Again, female CON-V rats (392ms, SEM = 35.9) were faster to collect the food reward than CON-V and CON-LM males (485ms, SEM = 62.7). At the 25ms cue duration, no factors affected latency to collect reward (all p’s>0.20). Lastly, latencies to collect reward during non-cue trials were examined for differences across groups. Males and females significantly differed in reward latencies on non-cue trials (F(1,66) = 5.93; p = 0.017): Females (370.4ms, SEM = 28.7) were quicker to collect the food reward at the end of correct trials than males (521.9ms, SEM = 61.8). No other factors affected reward collection latencies.

### 2.12 SAT Task

SAT scores were calculated following completion of the SAT task and analyzed as a function of cue duration (See Figure 5ABC). Again, a repeated measures ANOVA was used to determine if performance on the SAT declined as the cue duration. Similar to pSAT, sphericity was found to have been violated and after applying a Greenhouse Geisser correct, SAT performance significantly declined as a cue duration decreased (F(1.53, = 284.91, p < 0.0001). For the 500ms, 50ms, and 25ms cue duration, no significant main effects of Treatment, Drug, or Sex were detected, and no significant interactions were observed all (p’s > 0.05). Similarly, no significant differences were detected on measures of perceptual sensitivity, response bias, or the number of sessions required to master the task (all p’s > 0.05; Figure 5D-F).

**Figure 5:**
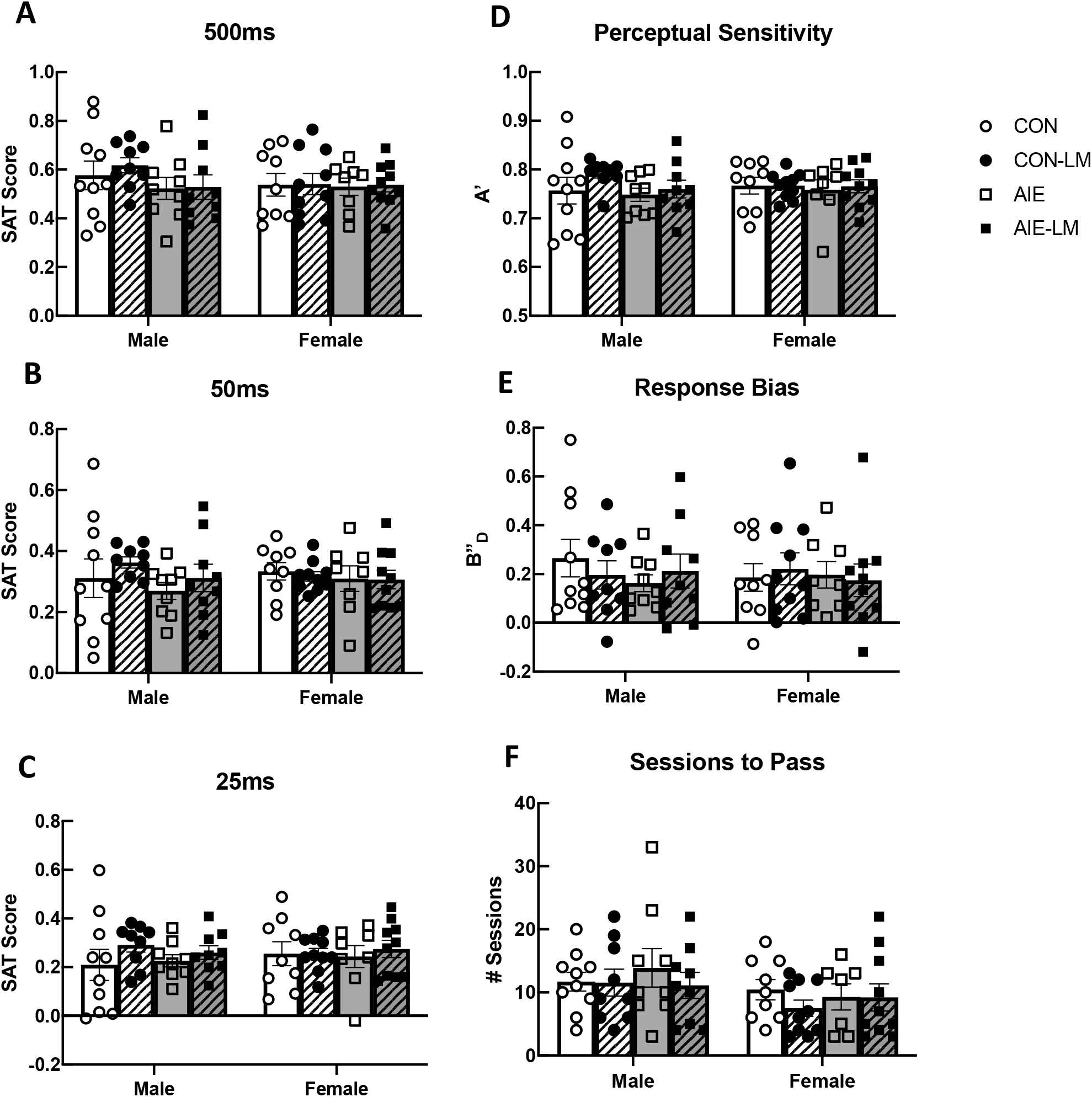
SAT Performance. SAT performance and Signal Detection Theory measures during the house light on condition. Data represent group means ±SEM. In both male and female rats, no significant group differences were found in SAT scores across the 500ms, 50ms, and 25ms cue durations (**A-C**). Groups did not differ in Signal Detection Theory measures of Perceptual Sensitivity or Response Bias, nor were there differences in the number of sessions required to complete the SAT task (**D-F**).

Latencies to lever press and reward were also examined. While no significant differences in latency to lever press were detected in the 500ms, 50ms, 25ms, and the non-cued trials (all p’s > 0.05), specific groups did not differ significantly in the latency to collect reward (all p’s > 0.05).

### 2.13 mPFC GRAB ACh 3.0 During SAT

ACh signaling was measured through in vivo-fiber photometry during the sustained attention task, trials were broken down to record responses to cue presentation, as well as response to reward (See Figure 6A-H for group averaged recordings). Area under the curve measurements of ACh activity in the mPFC did not reveal significant differences during cue presentation of trials of hits or misses (all p’s > 0.05; Fig 6IJ). Following Hits, a significant main effect of Treatment was observed (F(1,61) = 5.96, p = 0.017), where AIE treated rats had a lower AUC than CON rats (Figure 6K). In addition, a Treatment X Drug interaction was also observed following hits (F(1, 61) = 7.04, p = 0.01). Post hoc testing revealed that CON-V rats had a greater AUC than CON-LM (p = 0.036), and AIE-V treated rats, but not AIE-LM rats. Significant differences in AUC during misses were not observed across Treatment or Drug exposure (all p’s > 0.05, Figure 6L). Following correct rejections during non-signal trials, a main effect of Treatment was observed (F(1,61) = 4.74, p = 0.033), where AIE treated rats had a lower AUC compared to CON rats (Figure 6M). A significant Treatment x Drug interaction was also detected during correct rejections (F(1,61) = 4.58, p = 0.036). Again, post hoc testing revealed that AIE treated rats had a lower AUC following the lever press than CON rats. Lastly, during false alarm responses on non-signal trials, a significant Treatment x Drug interaction was observed (F(1,61) = 4.59, p = 0.036). Post hoc tests revealed that AIE treated rats had a lower AUC during the post response period compared to CON rats (p = 0.024, Figure 6N).

**Figure 6:**
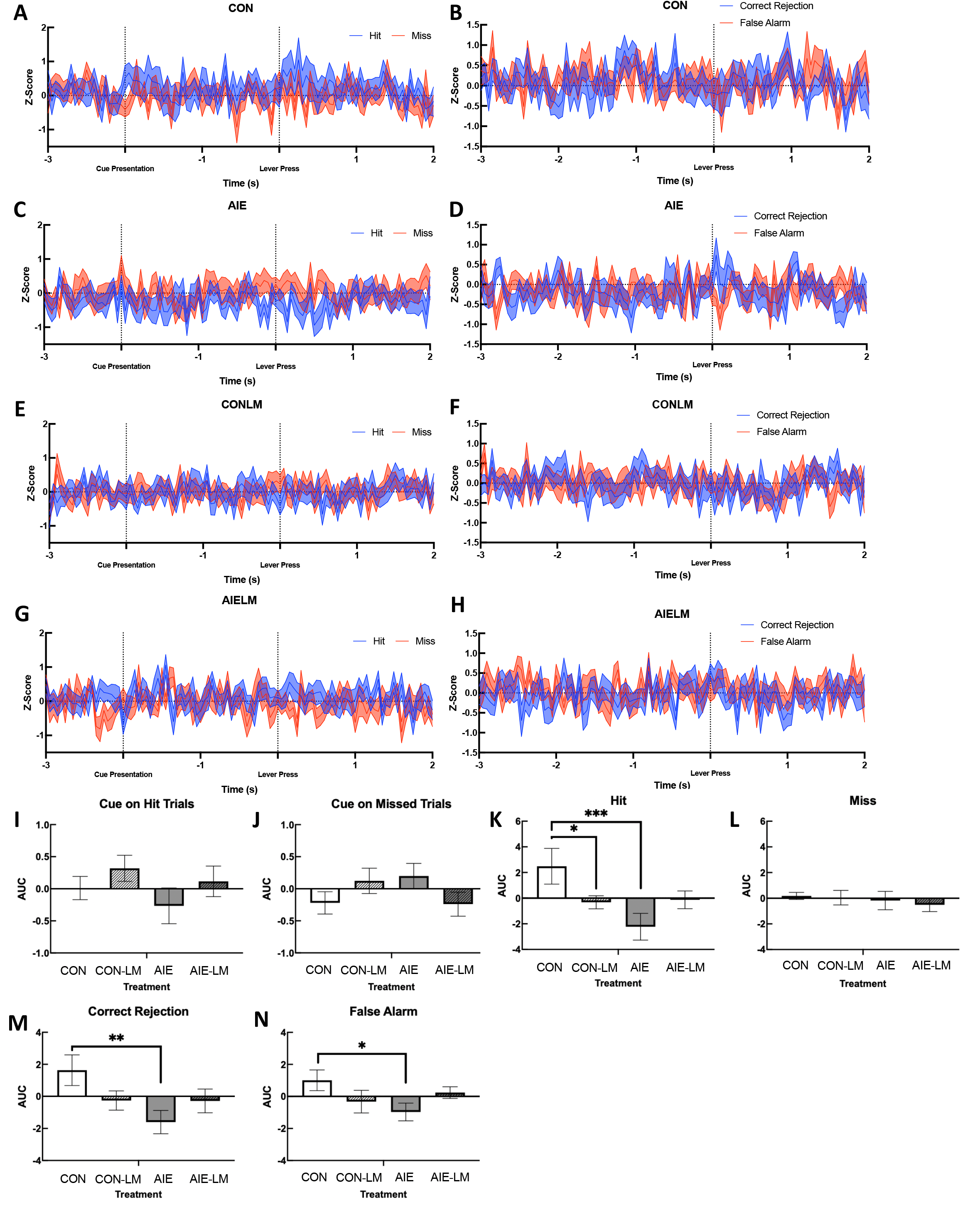
ACh 3.0 Activity During SAT and AUC measures. Example of a plot of group averaged Grab ACh 3.0 activity recorded via fiber photometry during signal (**A**, **C**, **E**, **G**) and non-signal trials (**B**, **D**, **F**, **H**) of SAT. Data represents Z-score transformation of signal relative to baseline recording during the previous intertrial interval. Area under the curve measurements during cue presentation, response selection, and the post reinforcement period during the SAT task (**I-N**). Data represent the AUC ±SEM. No group differences were observed during the cue presentation on hit and miss trials (**I-J**). However, CON-V rats had a greater AUC during the post response period following hits compared to AIE-V and CON-LM treated animals. LM11-31A treatment prevented this defict (**K**). No group differences were present in the AUC during misses (**L**). CON rats also had greater AUC compared to AIE rats in the post response period following Correct Rejections (**M**) and False Alarms (**N**), which were not corrected by LM11-31A treatment in AIE treated rats. * Indicates p < 0.05; ** p < 0.01, *** p 0.005.

When examining ACh signaling during cue presentation, it was found that groups did not differ in peak Z-score recorded at the 500ms cue duration in trials that would ultimately become hits (all p’s > 0.05; Figure 7A) or misses (all p’s > 0.05; Figure 7B). The peak z score in response to the 50ms cue during trials that would become hits, did not differ across Treatment or Drug conditions (all p’s > 0.05; Figure 7C). During cue presentation of 50ms miss trials, a significant Treatment X Drug interaction was identified (F(1,61) = 7.21, p = 0.0093; Figure 7D). During 50ms cue trials, ACh activity was significantly lower in AIE-V treated male and females compared to CON-V (p = 0.0098) and AIE-LM (p = 0.029) conditions. During 25ms trials, a significant Treatment effect was observed on hit trials (F(1,61) = 5.23, p = 0.025; Figure 7E), but not miss trials (p’s = 0.98; Figure 7F): CON-V and CON-LM treatment conditions had greater peak z scores than AIE-V and AIE-LM treatment conditions. Main effects of Drug were not found on 25ms hit trials (p = 0.82) or miss trials (p = 0.31), nor were there interaction effects on hit trials (p = 0.92) or miss trials (p = 0.56).

**Figure 7:**
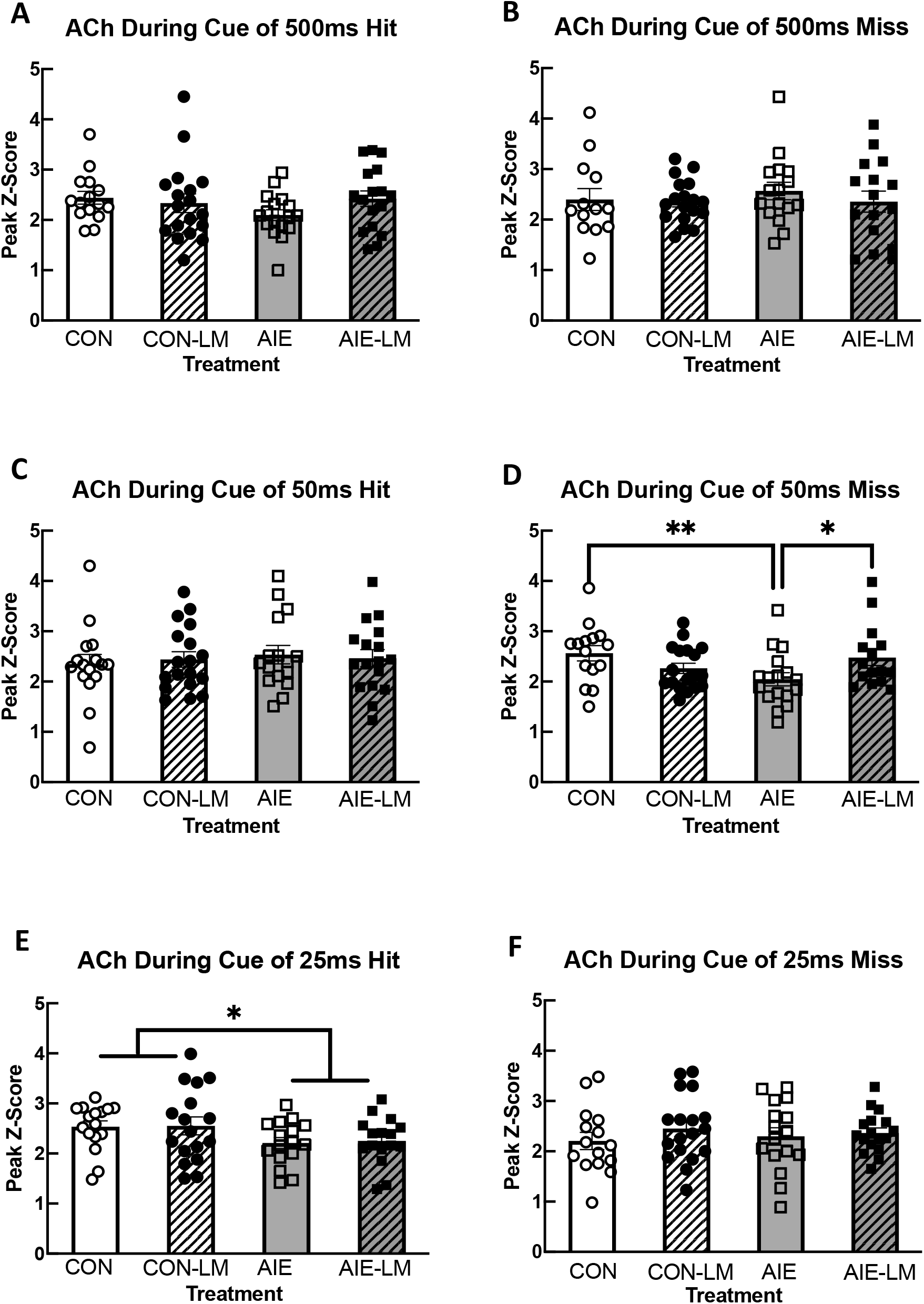
GRAB ACh 3.0 Activity During Cue Presentation. Grab ACh 3.0 recording during cue presentation on the SAT task. Data represents mean peak Z-score ±SEM. No group differences were found in ACh 3.0 activity during cue presentation on correct 500ms, 50ms, or 25ms cue duration trials (**A, C, E**). No group differences were found in peak z-score during cue presentation of 500ms cue miss trials (**B**). However, during 50ms cue presentation of miss trials, AIE-V treated animals had a significantly lower peak z-score relative to CON-V and AIE-LM treated animals (**D**). No group differences were observed on the 25ms cue trials that resulted in a miss (**F**). * Indicates p < 0.05; ** p < 0.01.

ACh activity was also measured in the form of peak z score immediately following hits, misses, correct rejections, and false alarms. When examining cued trials, it was found that ACh activity did not differ on 500ms trials following hits (all p’s >0.09; Figure 8A) or misses (all p’s; Figure 8B). During the reward period of 50ms hits, a significant Treatment X Drug interaction was detected (F(1,63) = 9.34, p = 0.0033; Figure 8C). Following correct choice on 50ms trials, the peak z score of ACh activity was significantly lower in AIE-V rats when compared to CON-V (p = 0.021) and AIE-LM treated animals (p = 0.017). A significant Treatment X Drug interaction was also observed following misses on 50ms trials (F(1,61) = 10.53, p = 0.0019; Figure 8D). The peak z score of ACh activity in the mPFC of CON-V rats were significantly higher than AIE-V and CON-LM groups (both p’s <0.04). Moreover, the AIE-LM condition had higher ACh peak z-score than AIE-V (p = 0.02) rats. No significant differences were observed in the period of recording following hits and misses on the 25ms cue duration (all p’s > 0.05; Figures 8E, F). In addition, no group differences were observed in the peak z score of ACh activity on the non-trials, following correction rejections (all p’s > 0.05; Figure 8G) and false alarm trials (all p’s > 0.05; Figure 8H).

**Figure 8:**
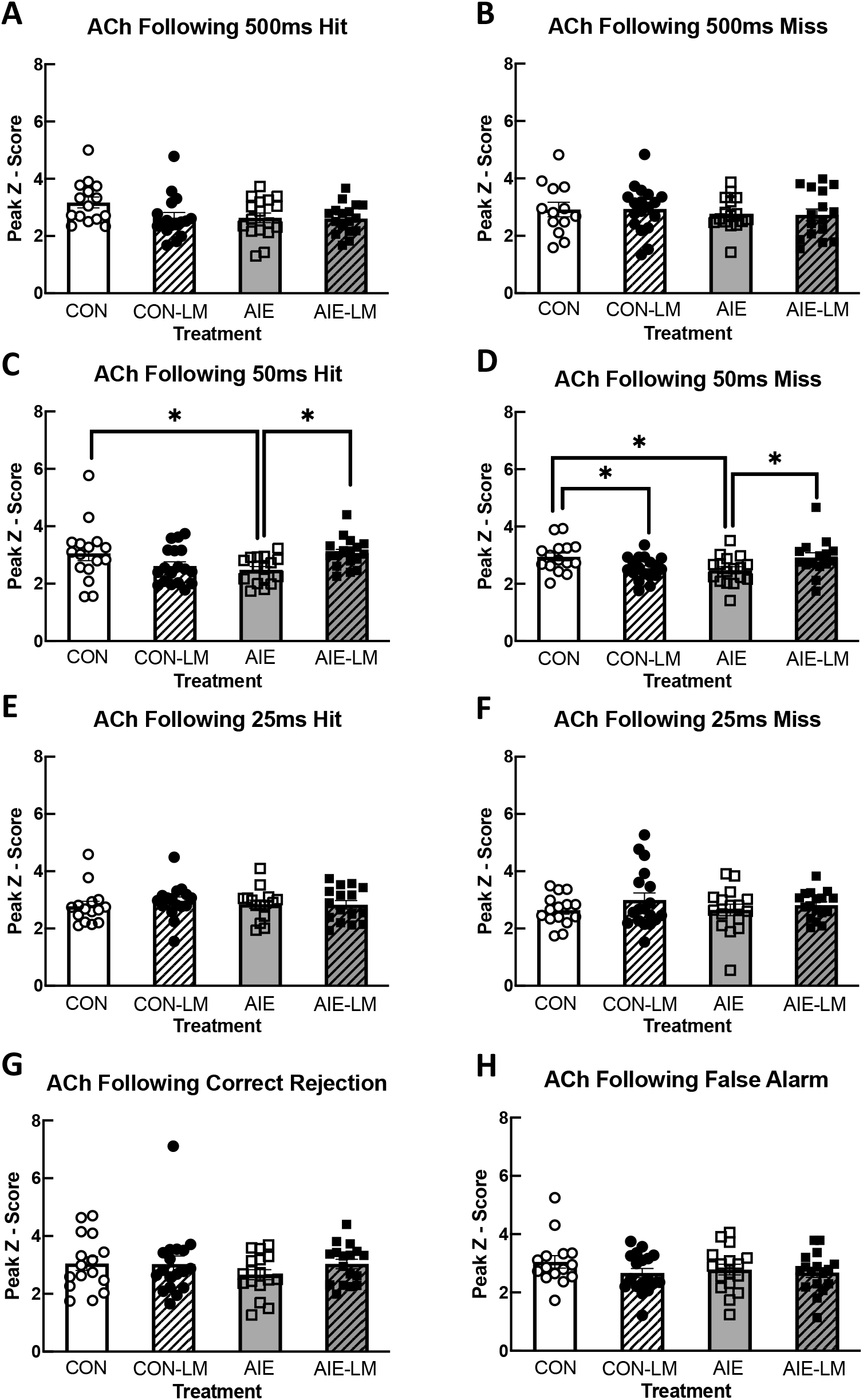
GRAB ACh 3.0 recording during Hits, Misses, Correct Rejections, and False Alarms. Grab ACh 3.0 activity measured with fiber photometry during the post response period of correct and incorrect trials (**A-F**). Data represent the mean peak z-score ±SEM. While no group differences were evident following 500ms cue trials (**A-B**), the peak z-score of ACh 3.0 activity was significantly lower in AIE-V treated rats compared to CON-V and AIE-LM rats following Hits and Misses on 50ms trials (**C-D**). Following a miss on the 50ms cue trial, peak z-score of GRAB ACh 3.0 was lower in the CON-LM treatment condition compared to CON-V. Groups did not differ in peak z-score following 25ms Hits or Misses (**E-F**). Groups also did not differ in peak z-score following correct rejections and false alarms of nonsignal trials (**G-H**). * Indicates p < 0.05.

500ms, 50ms, and 25ms cue duration trials were averaged together to determine if groups differed in peak z score of cued trials in general. Examining the peak z score of ACh activity during the cue of hits revealed a Treatment X Drug interaction (F(1,64) = 8.11, p = 0.0059; Figure 9A). Fisher’s LSD post hoc analyses revealed that AIE-V rats had lower peak ACh release than CON-V (p = 0.0005), AIE-LM (p = 0.0083), and CON-LM (p = 0.017) rats. It was also found that ACh signal significantly differed in the post response period of hit trials as a function of Treatment X Drug (F(1,64) = 4.33; p = 0.041; Figure 9C). Fisher’s LSD revealed that CON-V treated animals had greater peak z scores than AIE-LM (p = 0.015) and CON-LM (p = 0.028) groups. Interestingly, groups did not differ in response to the cue of miss trials, or in the post response period following a miss (all p’s > 0.05; Figure 8BD).

**Figure 9:**
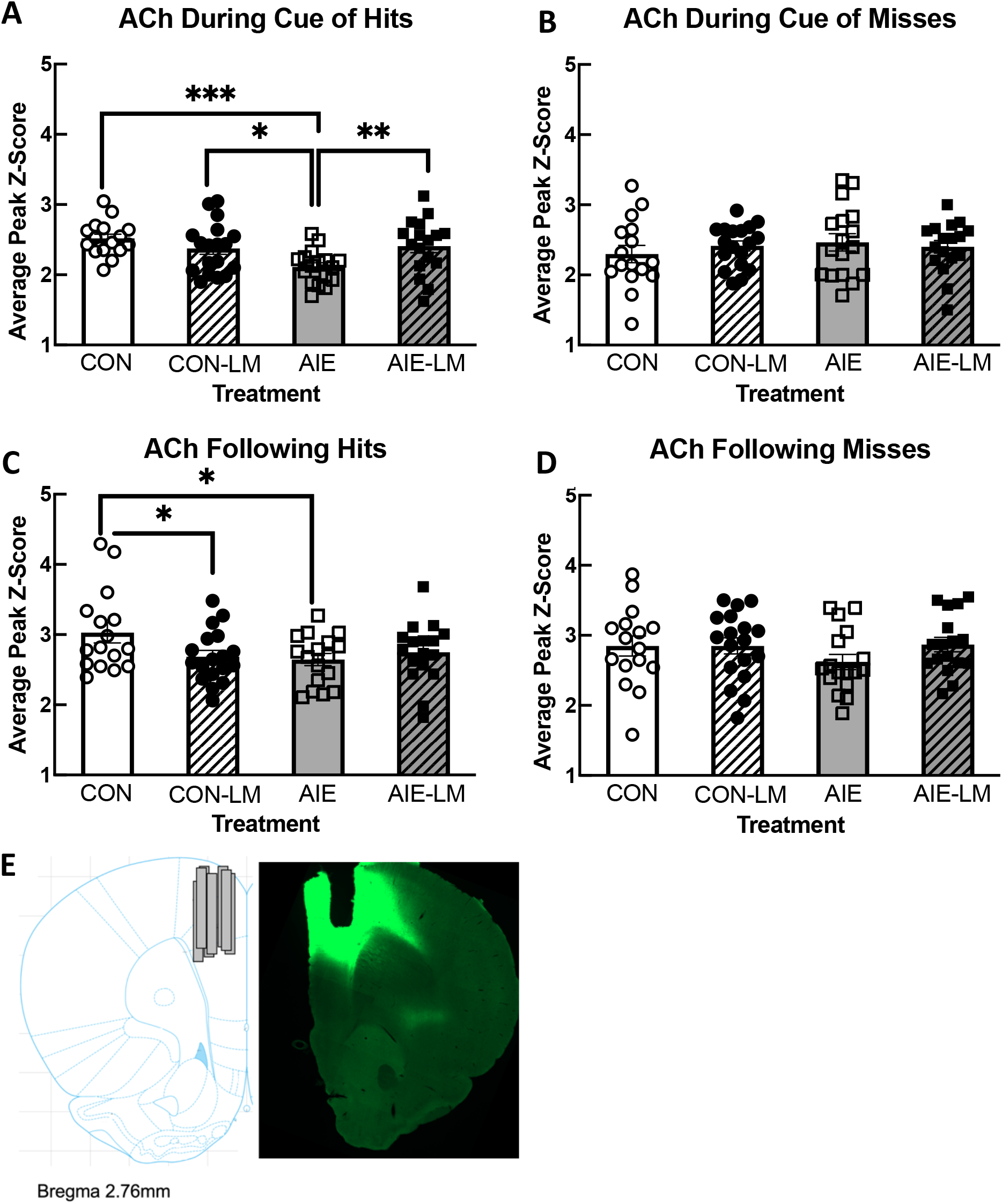
GRAB ACh 3.0 Activity Collapsed Across Cue Duration. Average Grab ACh 3.0 activity in the mPFC during cue presentation and the post response period across all cue durations. Data represent the group mean peak z-score ±SEM. During cue presentation on hit trials, AIE-V treated animals had significantly lower peak z-score of ACh 3.0 activity compared to CON-V, CON-LM, and AIE-LM treatment conditions (**A**). During the post response period of hit trials, CON-V treated animals had higher peak z-score ACh 3.0 activity compared to CON-LM and AIE-V treatment groups (**B**). Group differences in peak z-score were not detected during cue presentation of miss trials (**C**), or during the post response period of miss trials (**D**). * Indicates p < 0.05; ** p < 0.01; *** p < 0.005. Sample Grab ACh 3.0 viral expression and fiber optic cannula placement in the mPFC (2.70 AP, +/- 0.7ML, -3.0 DV); Image from Paxinos and Watson (2014) (**E**).

### 2.14 NbM ChAT+ TrkA+ Cell Counts

ChAT-TrkA co-expression was found to be significantly affected by Sex as males had more ChAT+TrkA+ cells than females (F(1, 66) = 4.22; p = 0.043; Figure 10A). Since males and females differed in ChAT+TrkA+ expression, separate analyses were run for males and females. In males, a Treatment X Drug interaction was found (F(1,34) = 17. 37, p = 0.0002). AIE led to a reduction in the number of ChAT+TrkA+ neurons in the NbM in males compared to CON males (p = 0.0006). However, LM11A-31 recovered the ChAT+TrkA+ population in AIE rats (p <0.0001). In females, a significant Treatment X Drug interaction was detected (F(1,33) = 7.82; p = 0.008): AIE females had fewer ChAT+TrkA+ cells than CON females (p = 0.015). AIE-LM females also had more ChAT+TrkA+ cells in the NbM compared to AIE-V females (p = 0.055).

**Figure 10.**
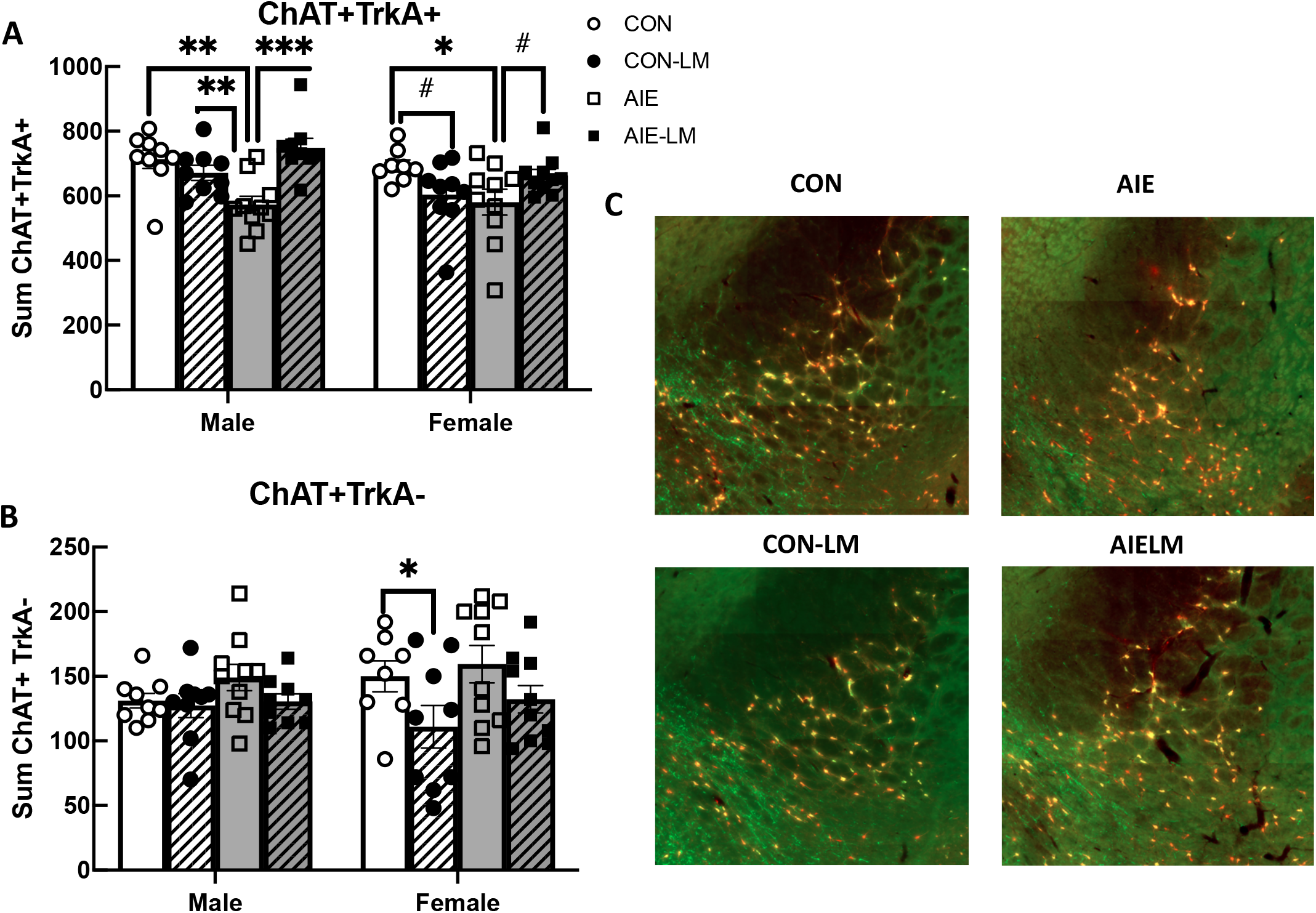
ChAT+, TrkA+ NbM Cell Counts. The sum number of fluorescently labeled ChAT+ neurons in the NbM and the number of ChAT+TrkA+ and ChAT+TrkA-phenotypes. Data represent group means of the total number of counted cells across four sections ±SEM. Group differences were evident in the ChAT+TrkA+ phenotype of cholinergic neurons in the NbM. In males, AIE-V treated rats had significantly fewer ChAT+TrkA+ labeled neurons than CON-V, CON-LM, and AIE-LM rats. In females, AIE-V treated rats also had fewer ChAT+TrkA+ cells than CON-V, CON-LM, and AIE-LM females, while CON-LM treated rats had fewer ChAT+TrkA+ cells than CON-V females (**A**). Group differences in ChAT+TrkA-cell counts in males were not evident; however, CON-LM females had fewer ChAT+TrkA-cells in the NbM compared to CON-V females (**B**). Sample images of NbM sections for ChAT+TrkA+ (Gold) and ChAT+TrkA-(Red) (**C**). ChAT-TrkA+ cells were not detected in these sections. All images were recorded at 20x magnification. * Indicates p < 0.05; ** p < 0.01; *** p < 0.005; # indicates trending effect.

When examining the ChAT+TrkA-phenotype, differences were not observed as a function of Treatment (p = 0.11), or Sex (p = 0.65); however, LM11A-31 did have an effect on the number of ChAT+TrkA-cells in the NbM (F(1,31) = 7.36, p = 0.011; Figure 10B), driven by reductions in CON-LM females compared to CON-V females.

## Discussion

There are several hypotheses about the neural dysfunction associated with adolescent heavy alcohol exposure [2, 46, 47] Recently, neurotrophin disruption has emerged as a key player in alcohol related brain damage—particularly loss of the cholinergic phenotype following adolescent binge type ethanol exposure [22, 48]. Although modulating the p75NTR has been shown to be neuroprotective, especially in basal forebrain cholinergic cell populations [33, 36, 37], no previous work has been done to determine if targeting the p75NTR during binge ethanol exposure would protect cholinergic neuronal phenotypes and or improve long-term neurocognitive outcomes. Moreover, no work has been done to investigate the developmental consequences of p75NTR manipulation or whether disrupting this signaling during critical periods of development lead to aberrant changes in brain and behavior. Here we show that LM11A-31 treatment during AIE protects attentional performance, cortical ACh activity, and basal forebrain cholinergic cell populations, regardless of sex. However, treatment with LM11A-31 in control adolescent female rats suppressed some NbM cholinergic cell populations. These results suggest that blocking the p75NTR in control animals may produce adverse consequences in cholinergic phenotype expression.

The first goal of this study was to characterize the attentional performance of rats that had underwent AIE treatment. Given that AIE has been consistently shown to lead to loss of basal forebrain cholinergic phenotype expression, and reductions in cortical ACh release [19–22, 25, 49], we expected that AIE would also negatively affect attentional performance, as ACh is a key driver of attention [6, 8, 50]. We found that AIE caused an impairment during SAT pretraining where AIE treated animals displayed attention deficits at each stimulus duration. However, co-treatment of LM11A-31 during AIE blocked these deficits from occurring. Through the application of signal detection theory, we are also able to parse out more discrete cognitive operations on this task through measurements of perceptual sensitivity (A’) and response bias (B”). Perceptual sensitivity measures bottom-up or exogenous attentional performance, in that is driven by the intensity of environmental stimuli relative to background noise [39, 41, 51, 52]. Bottom-up attention is also known as stimulus-driven attention, as it is the automatic capture of attention by external stimuli that stand out due to their distinct features, such as color, motion, or intensity. In contrast, response bias is a measure of top-down attentional control whereby selective internal processing, by factors such as goals, expectations, and previous experience of cues facilitates a greater signal to noise ratio of stimulus detection through changes in response strategy [39, 41, 51, 52]. We found AIE-induced impairments in both perceptual sensitivity and response bias during pSAT, but not SAT. It should be noted that changes in response bias, do not produce changes in perceptual sensitivity, or vice versa [41].

It appears that AIE treatment results in the failure to adopt a conservative approach to cue detection, evident with B” being closer to 0. AIE treated rats, along with the CON-LM group were more likely to indicate the presence of the cue, regardless of whether the cue is presented when compared to CON and AIELM treatment groups. Response bias is reflective of top down attentional control [39, 52, 53], and choline transporter mutations (Val89) coincide with reduced attentional performance on SAT, reductions in cholinergic signaling, and a more liberal response bias compared to WT animals [54]. During sustained attention, intact animals develop a conservative response bias to cue detection, where top-down attentional control on this task generally favors a bias in indicating the absence of the cue [39, 54]. Response bias is influenced by the probability of cue presentation, and as the frequency of cue presentation increases, response bias shifts towards more liberal indication that the cue is presented [55]. Cue presentation at the 25ms and 50ms duration are often interpreted as never having been present (i.e. perceived decrease in frequency of cue presentation); thus, response bias generally shifts towards indicating that the cue was not presented, reflected by an increase in B” values. This lack of response bias is in AIE treated animals may be indicative of an impaired top-down attentional control during pSAT. In addition, reductions in A’ in rats exposed to AIE, relative to controls, suggest that AIE decreases cue detection ability. Both decrements in response bias and perceptual sensitivity were prevented through LM11A-31 modulation during AIE treatment. However, treatment with LM11A-31 during the adolescent period in control rats also inhibited top-down attentional control, as measured by decrements in B”. This suggests that blocking the p75NTR during adolescence may have had adverse developmental outcomes in the ability of cholinergic neurons to mediate top-down attentional control. In conjunction with the findings that LM11A-31 treatment in female control rats led to a loss of cholinergic neurons in the NbM, indicate that LM11A-31 may contribute to cholinergic degeneration and attentional deficits in intact animals.

Performance on the pSAT and SAT tasks also revealed a few interesting sex-specific effects. On average, female rats took longer to master the pSAT task and demonstrated increased latencies to lever press at shorter cue durations than male rats, but not during 500ms cue or no cue trials. While findings of sex differences in attention are mixed, males have been found to have greater vigilance, while females tend to have reductions in reaction time-but greater inhibitory control [39, 56–62]. Greater inhibitory control typically observed in females most likely contributed to the greater response bias observed in female rats compared to male rats [56, 57, 61]. In addition, female rats also demonstrated greater bottom-up attention than male rats, as revealed through higher perceptual sensitivity scores on pSAT-which conflicts with previous findings in females [39, 62, 63]. Collectively, the differences between male and female rats in perceptual sensitivity and response bias also led to females having higher SAT scores, at each cue duration, compared to males.

Although differences between AIE and water treated animals in sustained attention performance were not evident under the more taxing condition when the house light was on. The SAT was developed to assess sustained attention which has been demonstrated to elicit greater ACh release (150% increase) compared to retractable lever or FI-9 tasks (50% increase) [64]. Moreover, optogenetic stimulation of NbM cholinergic neurons during cue presentation significantly improves attentional performance, while inhibition of NbM cholinergic neurons impairs cue detection on this task [50]. It is possible that attention deficits in AIE treated animals may have been masked by increases in task difficulty—or a floor effect. We saw a drop in performance in all rats when rats were transferred to the SAT version, which is expected [39].

Previous research has shown that ChAT phenotype suppression in the basal forebrain, induced by IgG Saporin lesions, leads to impairments on SAT [6]. The lack of and AIE effect on the more complex SAT task could also be due to the fact that AIE does not produce the same magnitude of cholinergic pathology (25% loss) typically observed following cholinergic lesion studies (∼50% loss; [6, 65]. Although sustained attention has not been examined in developmental models of ethanol exposure, studies in human adolescents have shown that a history of alcohol misuse is associated with worsening performance on sustained attention [66–68]. The findings from this study can extend the observed sustained attention deficits in human adolescents, to rodents treated with the adolescent intermittent ethanol paradigm.

In animal models of adolescent binge drinking, previous work using the Five choice serial reaction time task (5-CSRTT) and attentional set shifting reveal mixed results pertaining to attentional deficits [24, 69–73]. Previous work investigating differences in attentional performance following 2 g/kg ethanol exposure on a two day on two day off cycle from in early adolescence (PND 30-45) in D2 and B6 male mice found that ethanol did not produce attentional impairments on the 5-CSRTT, while late adolescent to early adulthood treatment (PND 45-60) increased the number of premature responses [69]. Conversely, 5g/kg ethanol exposure on a two-day-on two-day-off schedule from PND 25-57 in male Wistar rats had improved attentional performance on the 5-CSRTT, stemming from fewer omissions, than control rats [70]. Similarly, 5g/kg ethanol treatment from PND 25-54 in Sprague Dawley rats has been shown to increase [72] and decrease attentional set shifting performance [24, 55, 73]. The conflicting findings on attentional performance following AIE could be due to underlying differences in other cognitive domains that affect performance on 5-CSRTT and ASST, namely impulsivity, cognitive flexibility, and behavioral flexibility.

Recording cholinergic activity in the mPFC via fiber photometry during SAT revealed that AIE treated animals have blunted ACh responses to cue presentation and reward, and that LM11A-31 treatment during ethanol exposure had a protective effect on prefrontal cortical ACh signaling. Previous experiments in cortical ACh signaling following AIE have identified blunting of activity related ACh release in the mPFC and the OFC through in vivo-microdialysis during a spatial working memory task [24, 25]. However, microdialysis is limited in its ability to record ACh in more temporally restricted behavioral epochs and is unable to disentangle tonic and phasic ACh release. Previous work has shown that optogenetic stimulation of basal forebrain cholinergic neurons during cue presentation, facilitates cue detection on SAT, while optogenetic inhibition of this cholinergic population inhibits SAT performance [50]. Moreover, through the use of choline sensitive microelectrodes, prefrontal cortical increases during cue detection[8] and cholinergic basal forebrain neuron activity increases in response to reward [74]. Interestingly, although AIE treated animals did not differ in behavioral indices from CON or LM11A-31 treated animals during the SAT task, differences in ACh activity, and loss of cholinergic neurons in the NbM persist. While reductions in PFC ACh activity were found in AIE treated animals despite similar performance on SAT, prefrontal cortical reciprocal projections to the posterior parietal cortex and basal forebrain likely compensate for the reductions in ACh during cue detection. Here, the PFC can mediated top down attentional control can through PFC to posterior parietal cortex glutamatergic transmission [75].

Results from our histological examination in NbM sections revealed that AIE produced reductions in ChAT+ expression in male and female rats, as shown in previous studies [21, 22, 24, 25, 76]. While we do observe differences in performance in pSAT between males and females, this difference most likely results from the aforementioned differences in approach behavior and not differences in underlying cholinergic circuits. Uniquely, we found that it was the co expression of ChAT+TrkA+ that was significantly reduced following treatment with AIE, but LM11A-31 treatment spared this population. Although the majority of NbM cholinergic neurons also co-express TrkA, a subpopulation of cholinergic neurons in this region, did not express TrkA which may confer increased vulnerability to degeneration in this subpopulation. By inhibiting the p75NTR during AIE, basal forebrain cholinergic phenotype, cortical ACh release, and attentional performance were spared from impairments. Together, this indicates that the degeneration of cholinergic neurons in this model is mediated by proneurotrophin signaling at the p75NTR. Previous work found that loss of Trk receptors precedes degeneration of cholinergic neurons, and that cholinergic cell death occurs through p75NTR only when Trk receptor expression levels fall below that of p75NTR [77]. This supports our findings that blocking p75NTR during ethanol exposure protects cholinergic cell populations. Interestingly, LM11A-31 also prevented the loss of the ChAT+TrkA+ subpopulation indicating that cholinergic access to cortically derived neurotrophins was also preserved, as Trk expression positively correlates with neurotrophin signaling [78–80]. In addition, it was observed that TrkA expression did not exist by without co-expression of ChAT in NbM sections. This finding suggests that the loss in cholinergic phenotype expression coincides with the loss in TrkA expression. Whether reductions in TrkA expression in NbM cholinergic neurons precede reductions in ChAT phenotype expression in AIE, have yet to be seen, although restoration with of the cholinergic phenotype also coincides with demethylation of cholinergic and TrkA promotors [49].

Interestingly, LM11A-31 treatment during adolescence in water-exposed females did lead to reductions in ChAT+, ChAT+TrkA+, and ChAT+TrkA-neurons. This, alongside reductions in mPFC ACh signaling during reward, and corresponding attentional impairments in CONLM rats suggest that inhibiting p75NTR activity during development in intact animals may invertedly produce a hypertrophic cholinergic state that ultimately leads to degeneration of the cholinergic system, during withdrawal from LM11A-31.

Taken together, these findings directly implicate AIE associated changes in p75NTR mediated signaling, in driving the loss of cholinergic phenotype in the basal forebrain, reductions in cortical ACh release, and behavioral impairments that persist despite discontinued exposure to ethanol in adulthood.

